# What evolutionary processes maintain MHCIIβ diversity within and among populations of stickleback?

**DOI:** 10.1101/2020.08.19.257568

**Authors:** Foen Peng, Kimberly M. Ballare, S. Hollis Woodard, Stijn den Haan, Daniel I. Bolnick

**Author notes:** **Correspondence:** Daniel I. Bolnick.

## Abstract

Major Histocompatibility Complex (MHC) genes encode for proteins that recognize foreign protein antigens to initiate T-cell mediated adaptive immune responses. They are often the most polymorphic genes in vertebrate genomes. How evolution maintains this diversity is still an unsettled issue. Three main hypotheses seek to explain the maintenance of MHC diversity by invoking pathogen-mediated selection: heterozygote advantage, frequency-dependent selection, and fluctuating selection across landscapes or through time. Here, we use a large-scale field parasite survey in a stickleback metapopulation to test predictions derived from each of these hypotheses. We identify over a thousand MHCIIβ alleles and find that many of them covary positively or negatively with parasite load, suggesting that these genes contribute to resistance or susceptibility. However, despite our large sample-size, we find no evidence for the widely-cited stabilizing selection on MHC heterozygosity, in which individuals with an intermediate number of MHC alleles have the lowest parasite burden. Nor do we observe a rare-allele advantage, or widespread fluctuating selection across populations. In contrast, we find that MHC diversity is best predicted by neutral genome-wide heterozygosity and between-population genomic divergence, suggesting neutral processes are important in shaping the pattern of metapopulation MHC diversity. Thus, although MHCIIβ is highly diverse and relevant to the type and intensity of macroparasite infection in these populations of stickleback, the main models of MHC evolution still provide little explanatory power in this system.

## Introduction

Genetic variation is the raw material for natural selection to act upon. Hence, there is a long history of evolutionary studies on how genetic variation is maintained in natural populations (Dobzhansky, 1982), either invoking selection or neutral processes. One of the most dramatic cases of genetic variation concerns polymorphism at Major Histocompatibility Complex (MHC) genes, which are commonly the most variable genes in vertebrate genomes (Sommer, 2005). Surprisingly, despite decades of research on MHC evolution, the evolutionary processes sustaining MHC polymorphism remain unclear, with inconsistent and often ambiguous support for competing hypotheses (Radwan, Babik, Kaufman, Lenz, & Winternitz, 2020). Here, we evaluate predictions of multiple competing MHC evolution models (adaptive and neutral), using an exceptionally large dataset of genetic variation and macroparasite infection in a metapopulation of threespine stickleback (*Gasterosteus aculeatus*) inhabiting lakes on Vancouver Island.

MHC genes play a key role in the adaptive immune system of vertebrates. The main function of these genes is to encode for cell surface proteins, which are used to detect foreign molecules. These molecules are then presented to T cells to initiate appropriate immune responses. The MHC gene family consists of two classes of genes, MHC I and MHC II, each of which can be represented by multiple paralog copies. MHC class I genes are expressed in all nucleated cells and typically bind to peptides derived from intracellular molecules, such as virus protein (Jensen, 2007). MHC class II genes (our focus here) are expressed only in antigen presenting cells, such as macrophages, dendritic cells, and B cells. These cells phagocytize extracellular material, digest the proteins, and if the MHC II proteins bind to the resulting fragments, these are presented on the cell surface to T cells, to possibly initiate an adaptive immune response (Jensen, 2007). The second exon of MHC II gene is often used to characterize the variability of MHC II genes, because it contains the antigen-binding domain (Sommer, 2005).

The maintenance of MHC diversity is often attributed to pathogen-mediated selection, although intraspecific processes, such as mate choice (Penn & Potts, 1999), are also potential sources of selection. As reviewed in Spurgin & Richardson (2010), there are three non-mutually exclusive pathogen-mediated selection mechanisms to explain MHC diversity: (1). *Heterozygote advantage*. Individuals with more MHC alleles might be able to recognize a more diverse set of parasite species, thus have fewer parasites. This advantage may lead to directional selection for ever-increasing diversity, but some widely-cited studies have presented evidence for intermediate optima (K. Mathias Wegner, Kalbe, Kurtz, Reusch, & Milinski, 2003). (2). *Negative frequency-dependent selection*. In a given population, common MHC alleles that protect against parasites impose selection on those parasites to change their antigens and evade recognition. Once parasites evolve strategies to evade these common alleles, individuals carrying the common alleles may be at a selective disadvantage. By contrast, a rare MHC allele imposes little selection on the parasites it recognizes (because it is rare), and so may tend to be more protective. For this negative frequency-dependence to work in the long run, it must on average hold that rare alleles are more protective (and hence more fit) than common alleles. (3) *Fluctuating selection*. Parasite community and parasite genotypes may vary in space and time. Given this, different host MHC alleles may be selectively favored in different locations and at different times. Gene flow between locations subject to divergent selection can sustain MHC diversity, as can temporally fluctuating selection.

Although many studies have examined the roles of the above-mentioned mechanisms in maintaining MHC diversity, the results are often mixed or even contradictory. For example, heterozygote advantage was supported in one experimental infection study with inbred mice (Penn, Damjanovich, & Potts, 2002), but the effect was not found in another study with outbred mice (Ilmonen et al., 2007). Theoretically, negative frequency dependent selection is more likely to explain the extraordinary diversity in MHC genes than heterozygote advantage (Borghans, Beltman, & De Boer, 2004). However, empirically demonstrating negative frequency dependent selection is more challenging, as the pattern is often also consistent with other mechanisms. For example, in house sparrows, population-specific MHC alleles were linked to host resistance or susceptibility to malaria (Bonneaud, Pérez-Tris, Federici, Chastel, & Sorci, 2006), but this pattern could be explained either by negative frequency dependent selection or spatially varying selection. A previous study of stickleback from three lake-stream population pairs (Stutz & Bolnick, 2017) tried to distinguish between negative frequency dependent selection or spatially varying selection. The study took advantage of migration between parapatric populations with different parasite communities, which in principle would allow them to partition benefits of rarity within populations, from costs of being a (rare) immigrant to a foreign habitat. Yet, no overall trend was found to support either mechanism. Such inconsistent or ambiguous results are typical in the MHC evolution literature. Therefore, a consensus regarding the relative importance of these mechanisms in maintaining MHC diversity has not been reached (Radwan et al., 2020).

In contrast to these adaptive hypotheses, whether and how MHC diversity is shaped by neutral processes is less studied. In small and/or isolated populations, MHC diversity is often heavily influenced by genetic drift, rather than balancing selection (Hedrick, Parker, & Lee, 2001; Miller & Lambert, 2004; Seddon & Ellegren, 2004). In large metapopulations, it is unclear what role neutral processes play in shaping MHC diversity.

MHC studies in threespine stickleback have generated some unique insights in the past (Eizaguirre, Lenz, Kalbe, & Milinski, 2012; Matthews, Harmon, M’Gonigle, Marchinko, & Schaschl, 2010; Mccairns, Bourget, & Bernatchez, 2011; Milinski et al., 2005; K. Mathias Wegner et al., 2003). For example, Wegner et al. (2003) conducted a study on 8 stickleback populations to examine the association between MHC diversity and 15 parasite species. They found populations exposed to a wider range of parasites had higher MHC allelic diversity, though MHC diversity also positively, albeit weakly, correlated with neutral diversity. Moreover, at the individual level, parasite burden was minimized in fish with an intermediate, rather than maximal, number of MHCIIβ alleles. This result supports the theoretical model of stabilizing selection on MHC heterogeneity (Nowak, Tarczy-Hornoch, & Austyn, 1992). The logic is that low MHC-diversity individuals do not have the capacity to recognize diverse parasites, but high MHC-diversity individuals deplete their T cells via negative selection on self-reactive T cells. The resulting paucity of T cell receptor diversity also limits their ability to recognize diverse parasites. However, this intermediate-advantage hypothesis, while widely cited, has not been replicated in many systems and remains controversial (Borghans, Noest, & De Boer, 2003). Another example of interesting insights from stickleback is the involvement of MHC alleles in mate recognition. It was found that female stickleback could choose their mate based on odor to optimize the number of MHC alleles in their offspring, though the molecular mechanism is unknown (Milinski et al, 2005).

In this study, we set out to test predictions associated with three main models of MHC polymorphism invoking parasite-mediated selection, using an extensive field survey of parasites in a metapopulation of threespine stickleback. We did not find evidence for stabilizing or positive selection for MHC heterozygosity, contrary to Wegner et al. (2003). Nor did we observe fluctuating selection across populations. Although we found significant associations between specific MHC alleles and parasite species both within and across populations, neutral processes in our dataset best explained within- and between-population MHC diversity in our data set.

## Materials and methods

### 1. Fish sampling and parasite identification

We used unbaited minnow traps to collect adult threespine stickleback from lake, river, and estuary sites on Vancouver Island in late May and early June 2009. Collections were approved by the University of Texas IACUC (07-032201) and a Scientific Fish Collection Permit from the Ministry of the Environment of British Columbia (NA07-32612). Fish were euthanized in MS-222 and then preserved in formalin, after the caudal fin of each individual fish was removed and preserved in pure ethanol for genotyping. Details are provided in Bolnick & Ballare (2020). These samples were used to examine within- and between-population variation in diet (Bolnick & Ballare, 2020), parasite community composition (Bolnick, Resetarits, Ballare, Stuart, & Stutz, 2019), and parasite species richness (Bolnick, Resetarits, Ballare, Stuart, & Stutz, 2020). The parasite infection data for this study are archived at Dryad Digital Repository (Bolnick and Ballare, 2020). Here, we focus on a random subset of 26 sampling sites (N = 1437 stickleback) for which we also genotyped individuals for MHC IIβ. For each stickleback individual, we recorded sex and body length and then dissected each fish to count and identify macroparasites as described in (Stutz & Bolnick, 2017).

### 2. Genotyping

The methods to sequence and genotype MHC IIβ alleles were identical to Stutz & Bolnick (2017). Briefly, genomic DNA was extracted from fin clips using a Promega Wizard 96-well extraction kit. We used PCR to amplify the second exon of MHC IIβ genes in each fish, with primers and PCR cycles as described in Stutz & Bolnick (2017). This exon contains the hypervariable peptide-binding region that binds to possible parasite antigens (Sommer, 2005). Each specimen was barcoded with a unique combination of forward and reverse primer tags for multiplexing. We used Quant-iT PicoGreen kits (Invitrogen P11496) to quantify DNA concentrations of magnetic bead-purified (Agencourt AMPure XP beads) PCR products, then pooled up to 400 samples in equimolar amounts to construct a library. We used Illumina Mi-Seq to sequence these multiplexed amplicon libraries. Then, we used a Stepwise Threshold Clustering (STC) program (Stutz & Bolnick, 2014), implemented in the AmpliSaS web software (URL: http://evobiolab.biol.amu.edu.pl/amplisat/index.php?amplisas; Sebastian, Herdegen, Migalska, & Radwan, 2016) to distinguish real sequence variants from sequencing error or PCR chimeras. The algorithm was originally validated by sequencing cloned amplicon products (Stutz & Bolnick, 2017). The software outputs a table of individual fish (rows) and unique MHC sequences (columns) with read depths.

To efficiently process data in AmpliSaS, we set the upper limit of read depth for each individual as 5000, which was sufficient to retrieve all the possible MHC alleles (Stutz & Bolnick, 2014). Because low sequencing coverage could bias the number of MHC alleles to be identified, individual fishes with coverage lower than 450 were excluded from this study (Fig. S1). After excluding the 160 individuals with low coverage (leaving N = 1277 individuals in the subsequent analyses), there was no longer a significant linear relationship between sequencing coverage and the number of MHC alleles (t = 1.46, p = 0.14). The number of unique MHC alleles was inferred based on the translated protein sequences, thus merging distinct exonic sequences that produce identical amino acid sequences.

### 3. Data analysis

#### a. Heterozygote advantage

We tested whether MHC diversity is associated with parasite burden at both the individual and the population level. At the individual level, we used parasite richness (*i*.*e*., the number of parasite species) to measure parasite burden. Given that MHC genes locate in many genomic loci and often act in a dominant manner in parasite resistance, we used allelic richness (*i*.*e*., the total number of unique MHC alleles found in an individual) in this study to evaluate the level of MHC heterozygosity. We used a mixed effect generalized linear model with Poisson distribution to evaluate the impact of MHC alleles on parasite richness. The fixed effects are the number of MHC alleles and log body length, which is known to affect parasite richness in stickleback (Bolnick et al., 2020) and in other fish species (Calhoun, McDevitt-Galles, & Johnson, 2018). We did not include sex as a main effect because we found no main effect of sex on parasite richness in a previous analysis of this dataset (Bolnick et al., 2020). Sample site was treated as a random effect. In addition, we also evaluated the optimizing hypothesis (*i*.*e*., MHC allelic richness is optimized at an intermediate level, driven by the negative selection pressure to decrease T cell depletion by self-recognition) by adding a quadratic MHC diversity term into the mixed effect model. We considered the interaction between site and the linear and quadratic effects of MHC, using a random slope effect across sites. Statistical support for the model terms was evaluated by AIC.

At the population level, we calculated the mean value of parasite richness and the average number of MHC alleles (per fish) in each site. We used a linear regression model to test if the mean value of parasite richness is associated with the average number of MHC alleles for each site.

#### b1. Frequency-dependent selection: is there a rare allele advantage?

If MHC alleles were under negative frequency dependent selection, then on average rare alleles must confer an advantage (in the form of lower parasite infection) compared to common alleles. In contrast, MHC alleles with high frequency would be less effective in parasite detection, because parasites would be evolving strategies to avoid detection by those alleles. We therefore expect a positive relationship between the frequency of an allele and its overall effect on parasite infection, where a negative effect denotes protection and a positive effect suggests susceptibility or survivor’s bias. Because MHC II genes scatter throughout multiple loci in fish genome (Kaufman, 2018), it is difficult to use allele frequency to estimate the abundance of MHC allele in the population. We used allele prevalence instead, which was calculated as the percentage of individuals carrying a focal allele in a population. Note that allele prevalence had different properties from allele frequency; for example, allele prevalence values of all the alleles in a population do not sum to 1. Similarly, we also calculated parasite prevalence in a population as the percentage of individuals infected by a focal parasite. Alleles and parasites that are too rare (<0.05 prevalence) or too common (>0.95 prevalence) do not provide sufficient variance to estimate effects. For every moderately prevalent MHC - parasite combination (both prevalence variable ranging from 0.05 to 0.95) in every sampling site, we used a generalized linear model with negative binomial distribution to test if the presence/absence of the focal MHC allele influenced the infection intensity of the focal parasite in that stickleback population. We corrected p values for multiple-comparison with BH method. The Z value of the regression models indicated the impact of the particular MHC allele on the infection rate by a particular parasite. We excluded the models disproportionately influenced by extreme values, *i*.*e*. the absolute Z value of a model would change over 0.5 if excluding the largest data point from the model. After iterating this procedure for all qualifying MHC-parasite combinations, we used another mixed-effect linear regression model to examine if the estimated effect sizes (Z) of MHC alleles were influenced by local allele prevalence. In this model, we treated both sampling site and focal parasite as random effects.

#### b2. Frequency-dependent selection: are allele effects inconsistent across lakes?

If stickleback and their parasites engage in a Red Queen race style co-evolution, the efficacy of any single allele will shift through time. For hosts and parasites with limited dispersal, physically disconnected sites are unlikely to be in the same phase of the arms race. As a result, a given MHC allele may have different effects on a given parasite from one site to the next: effective against defense in some places/times, ineffective or susceptible at others. Alternatively, if the same allele has similar effects on the same parasite across different lakes (without gene flow), such fluctuating frequency-dependent selection is unlikely. To test these alternatives, we first identified the moderately prevalent MHC-parasite combinations that were present in more than one site. For each qualified combination, we used a generalized linear model with negative binomial distribution to examine if the infection intensity of the parasite is influenced by the MHC allele, sampling site, and the interaction between site and MHC allele (this differs from the GLMs described in b1, which were done separately for each sample site). We corrected p values for multiple-comparison with BH method. We used the *anova* function in R to perform an analysis of deviance for each regression model. It reported the reductions in the residual deviance as each term of the formula was added in turn. We evaluated whether, across many parasite-allele combinations, more variation was explained by the focal allele’s main effect (implying consistent protection across populations), or allele × population interactions (inconsistent protection). Parasites transmitted by birds could spread in a larger spatial scale, so they are less likely to be engaged in evolutionary arms race. We did a Chi-squared test to test whether the parasite taxa, which have birds as final hosts, were more likely to be found in the models with significant main effect than in the models with significant allele x population effect.

#### c. Fluctuating selection

The parasite communities of stickleback differ significantly among the different sites on Vancouver Island (Bolnick et al., 2020; Stutz & Bolnick, 2017; Stutz, Schmerer, Coates, & Bolnick, 2015). This heterogeneity could select for different MHC genotypes at different sites. If it is true, we expect that the distributions of MHC genotypes and parasite species would be correlated across the stickleback metapopulation. The populations that have similar parasite communities should have similar MHC genotype compositions, and *vice versa*. To test this hypothesis, we first constructed a Euclidian distance matrix across all the sites for MHC alleles and for parasite species, respectively. Then we used Mantel test to examine if the two distance matrices were correlated. A previous analysis (Bolnick et al., 2019) found no significant effect of as-the-crow-flies (*i*.*e*., Euclidean distance) or as-the-fish-swims (*i*.*e*., the shortest in-water path) geographic distance on parasite community structure, so we omit that as a covariate in this distance analysis.

#### d. Neutral evolution

We previously generated reduced representation genomic sequence data for the populations in this study, using ddRADseq following the protocol of Peterson, Weber, Kay, Fisher, & Hoekstra (2012) modified as described in Stuart et al. (2017). From these data we estimated genome-wide mean heterozygosity for each individual fish, and from this population-level mean heterozygosity. We also calculated pairwise Weir-Cockerham adjusted F_ST_ between each pair of populations.

If MHC polymorphism is strongly affected by bottlenecks or other neutral population genetic processes, then we may expect that MHC diversity would be positively correlated with genomic mean heterozygosity. We tested whether the average number of MHC alleles (per fish) in each population varied as a function of the mean heterozygosity. We might then also expect that between-population MHC divergence is primarily a reflection of shared ancestry due to colonization processes or ongoing gene flow. To test this, we used a Mantel test to evaluate correlations between among-population MHC distance matrix and the F_ST_ matrix.

## Results

### Sequencing results and parasite information

We kept 1277 stickleback from 26 sites for subsequent analyses, after excluding 160 individuals whose genotype calls may have under-represented their actual diversity due to low read depth (<450 reads each; Figure S1). These 26 sites were from 7 different watersheds (sample site information is in Table S1). The habitat types included 21 lakes, two rivers, and three estuaries. Each sample site had an average sample size of about 50 fish, with a range from 18 to 78 (Mean = 49.11, SD=14.7). We identified a total of 1115 unique MHC alleles, 820 (73.54%) of which were private alleles found in only one population. On average, each fish had seven distinct MHC alleles, with a range from 3-15 (M=6.94, SD=1.87). 4.6% variation of MHC allelic richness could be explained by habitat type (ANOVA, habitat term, p < 2e-16), 1.7% variation could be explained by watershed (ANOVA, watershed term, p = 4.38e-05), and 23.0% of variation was explained by sample site (ANOVA, watershed/site term, p < 2e-16). Fish from river (M=8.38, SD=1.98) and estuary (M=7.1, SD=1.81) habitat had higher MHC allelic diversity than fish from lake habitat (M=6.77, SD=1.82).

In total, we identified 33 parasite taxa (parasite information is in Table S2). On average, each fish 2.5 different parasite taxa, with a range from 0 to 10 (Mean = 2.53, SD=1.74). Fish from river (M=1.94, SD=1.6) and estuary (M=1.15, SD=0.76) habitat had lower parasite burdens than fish from lake habitat (M=2.82, SD=1.74). 11.8% variation in parasite richness could be explained by habitat type, 4.9% variation could be explained by watershed (ANOVA, habitat term, p < 2e-16), while 28.3% of variation could be explained by sample site (ANOVA, watershed/site term, p < 2e-16). See Bolnick et al. (2019 and 2020) for further analysis of ecological factors structuring the parasite metacommunity (diversity, composition, co-occurrence) within and among lakes.

#### a. Heterozygote advantage

We found no evidence for an association between stickleback allelic diversity and parasite diversity, at either the scale of individual fish or among populations. At the individual level (Fig. 2A), the best mixed effect generalized linear regression model was the base model (AIC: 4226.02) that only included the fixed effect of log fish body length (coefficient = 1.24, z = 8.78, p < 2e-16) and the random intercept of sampling site (standard deviation: 0.47) conveying that populations differ in parasite diversity (as previously shown, (Bolnick et al., 2020)). Adding the fixed effect of MHC alleles (coefficient = 0.01, z = 1.22, p = 0.22) into the base model slightly decreased model fit (AIC: 4226.53). Adding the quadratic MHC fixed effect further decreased model fit (AIC: 4239.6). No additional explanatory power was given by site by MHC diversity interactions (linear or quadratic), so we have no support for the possibility that MHC diversity is under selection in some populations but not others (a summary of all the models described in this section is available in Table S3). This result suggested that MHC diversity did not have a significant impact on parasite richness.

**Figure 1.**
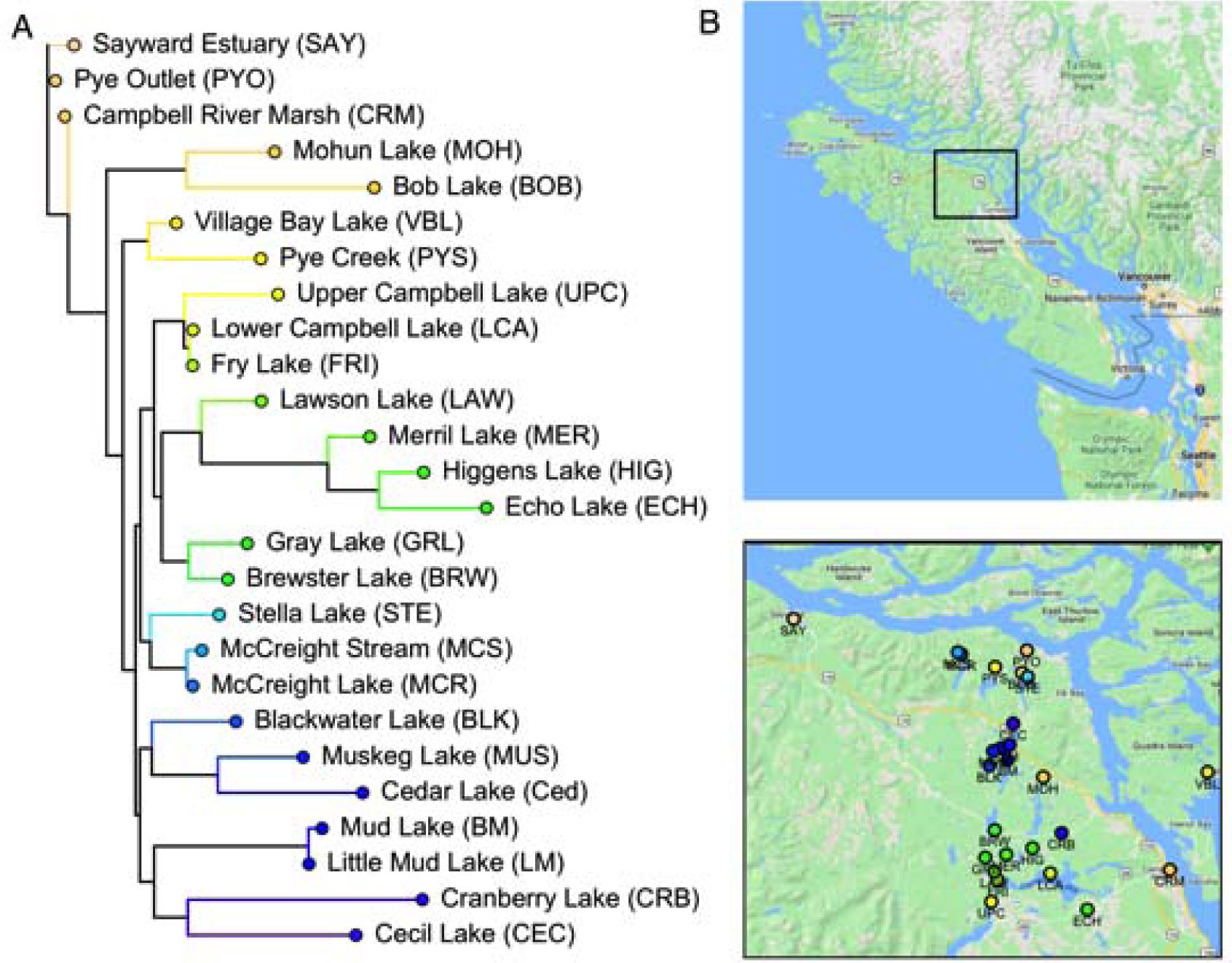
The phylogeny and geographic distribution of sampled stickleback populations. A. A rooted neighbor-joining tree of stickleback populations based on Fst calculated from SNPs obtained via ddRAD sequencing (Stuart et al. 2017). Site IDs are in parentheses. Note that Sayward Estuary (SAY), Pye Outlet (PYO), and Campbell River Marsh (CRM) are estuary habitat, and the rest are freshwater habitat. B. The distribution of sampling sites on Vancouver Island, Canada overlaid on Google Map. The lower panel is an enlarged view of the rectangle box in the upper panel. Each site is represented by a circle. The color of the circles is consistent with the tip label circles in A.

**Figure 2.**
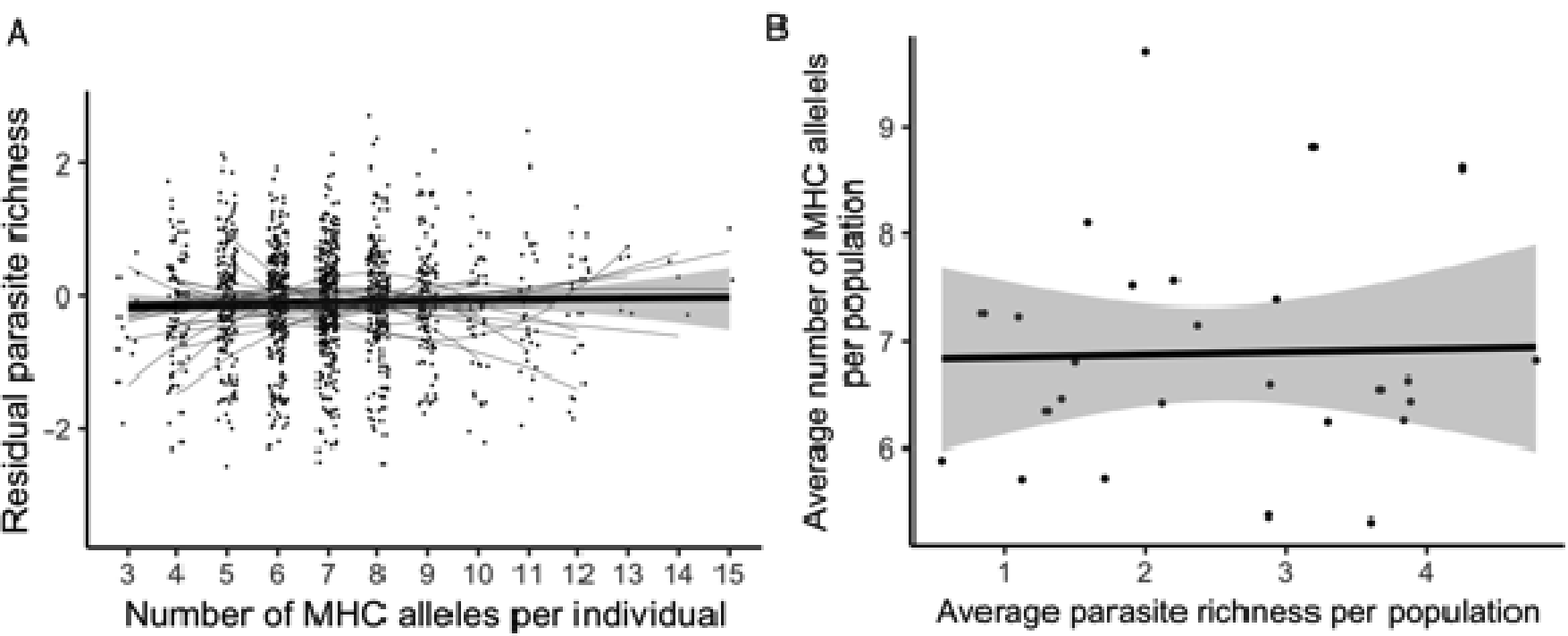
Testing predictions of heterozygote advantage. A. The relationship between the number of MHC alleles per individual and its residual parasite richness. For purposes of plotting, the residual parasite richness for each fish was calculated by using a mixed-effect generalized linear model to control for the random effect of sampling site and the fixed effect of the log body length (covariates in the statistical analysis we report in the text). Each dot represents a fish. The quadratic fit using either the full dataset or each population was represented by thick black line or thin grey lines, respectively. None of the quadratic fit using data from individual site was significant after correcting for multiple comparisons. B. The relationship between the average number of MHC alleles per-fish in a given population and the average per-fish parasite richness in that population. Each dot represents a population. The grey area represents 95% confidence interval. Note that x and y axes are reversed from panel A to B. In A we are specifically testing whether individual genotype affects their parasite burden, as the reverse direction of causation is not plausible. In B we are considering the hypothesis that existing parasite diversity is the driver of evolution of MHC diversity.

In the model just described, the random effect of sampling site explained a large portion of the total variance of parasite richness, as noted previously (Bolnick et al., 2020). Populations ranged from as few as an average of 0.5 macroparasite taxa per individual, to nearly 5 taxa per individual (Fig. 2B). We also observed significant variation in MHC allelic diversity between populations, ranging from as low as an average of 5.31 alleles per fish, to as high as 9.71 alleles per fish (Fig. 2B). Among populations, average MHC diversity was unrelated to average parasite richness (coefficient = 0.03, F = 0.02, df = 24, p = 0.89, Fig. 2B).

#### b1. Frequency-dependent selection: is there a rare allele advantage

Although we found many MHC-parasite associations, we found no tendency for alleles to be protective when rare, or vulnerable to infection when common. That is, the Z value of a given allele’s effect on a given parasite in a given site was independent of that alleles’ prevalence in that focal site. Across all sites, there were a total of 5623 MHC-parasite combinations that were moderately frequent and so included in the analysis. After excluding the models heavily influenced by outlier values with high leverage, we obtained 4130 linear regression models (see Fig. 3A for an example of this result from one population). 45 models had significant negative MHC-parasite associations and 45 models had significant positive MHC-parasite associations (see Fig. 3B for an example). Some MHC alleles, and parasites, are repeated across multiple regression models from different populations. None of the models were significant after we corrected for multiple-comparison with BH method. A mixed effect linear model showed that the impact of allele prevalence on Z values was not significant (coefficient = 0.09, t = 1.14, p = 0.27) (Fig. 3C). Therefore, this result did not provide evidence for rare allele advantage of MHC alleles on parasite infection intensities.

**Figure 3.**
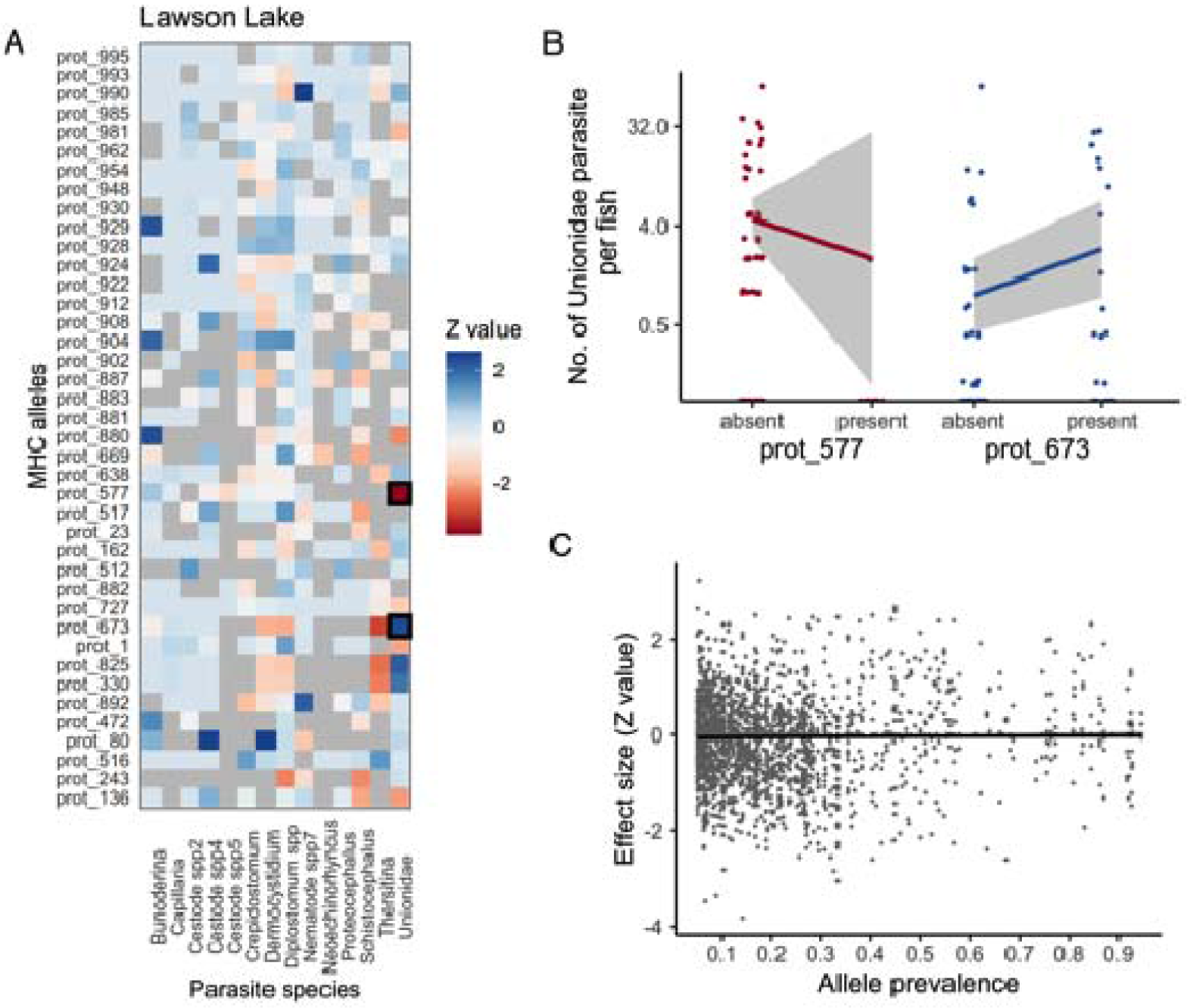
Testing predictions of frequency-dependent selection: rare allele advantage. A. The Z values of all MHC-parasite regressions in one sample site (Lawson Lake), as an example. Each model used data from one population and regressed the infection intensity of a focal parasite against the presence/absence of a focal MHC allele. Each row in the grid represents a MHC allele, and each column represents a parasite taxa. The color filling represents the effect size. The darker the blue color indicates a stronger positive association (having the allele confers higher infection load). The darker the orange color indicates a stronger negative association (e.g., resistance). The MHC-parasite regressions heavily biased by extreme values are filled as grey and excluded from further analysis. Two examples (highlighted in black boxes in A) are plotted in B. B. The infection intensity of parasite Unionidae is negatively and positively influenced by the presence/absence of MHC allele prot_577 and prot_673, respectively. Note the y axis of B is on log scale. C. The relationship between the allele prevalence of an MHC allele and its effect size (Z value) on the infection intensity of certain parasite. Each dot represents a MHC-parasite combination in a particular population. The solid line depicts the result of linear regression using all the data. The grey area surrounding the regression line indicates 95% confidence interval.

#### b2. Frequency-dependent selection: are allele effects inconsistent across lakes?

We found some cases where the allele effects are inconsistent across lakes (significant MHC allele × site interaction effects), suggesting there were potentially Red Queen style co-evolutionary arms race between stickleback population and parasites. We obtained 788 generalized linear models for each moderately prevalent MHC-parasite combinations which were present in more than one site. 53 models had a significant interaction term but an insignificant MHC term (11 of them involved parasites with birds as final hosts), suggesting inconsistent allelic effects across sites. For example, the allele prot_110 confers susceptibility (higher infection) in McCreight Stream, but has no effect in McCreight Lake (Fig. 4C). Although none of the models were significant after correction for multiple-comparison with BH method, the number of significant models was greater than null expectation with a significance level of 0.05 (792 * 0.05 = 39.6), so some of the model results were truly significant despite the issue of multiple comparison. 66 models had a significant MHC term but an insignificant interaction term (13 of them involved parasites with birds as final hosts) (Fig. 4B), suggesting that those alleles had consistent effects on the focal parasite across sites, and thus did not fit localized Red Queen race dynamics. Parasites with birds as final hosts were not more likely to be found in the models with significant MHC term than in the models with significant interaction term (Chi-squared test, X^2^ =0.004, p=0.95).

**Figure 4.**
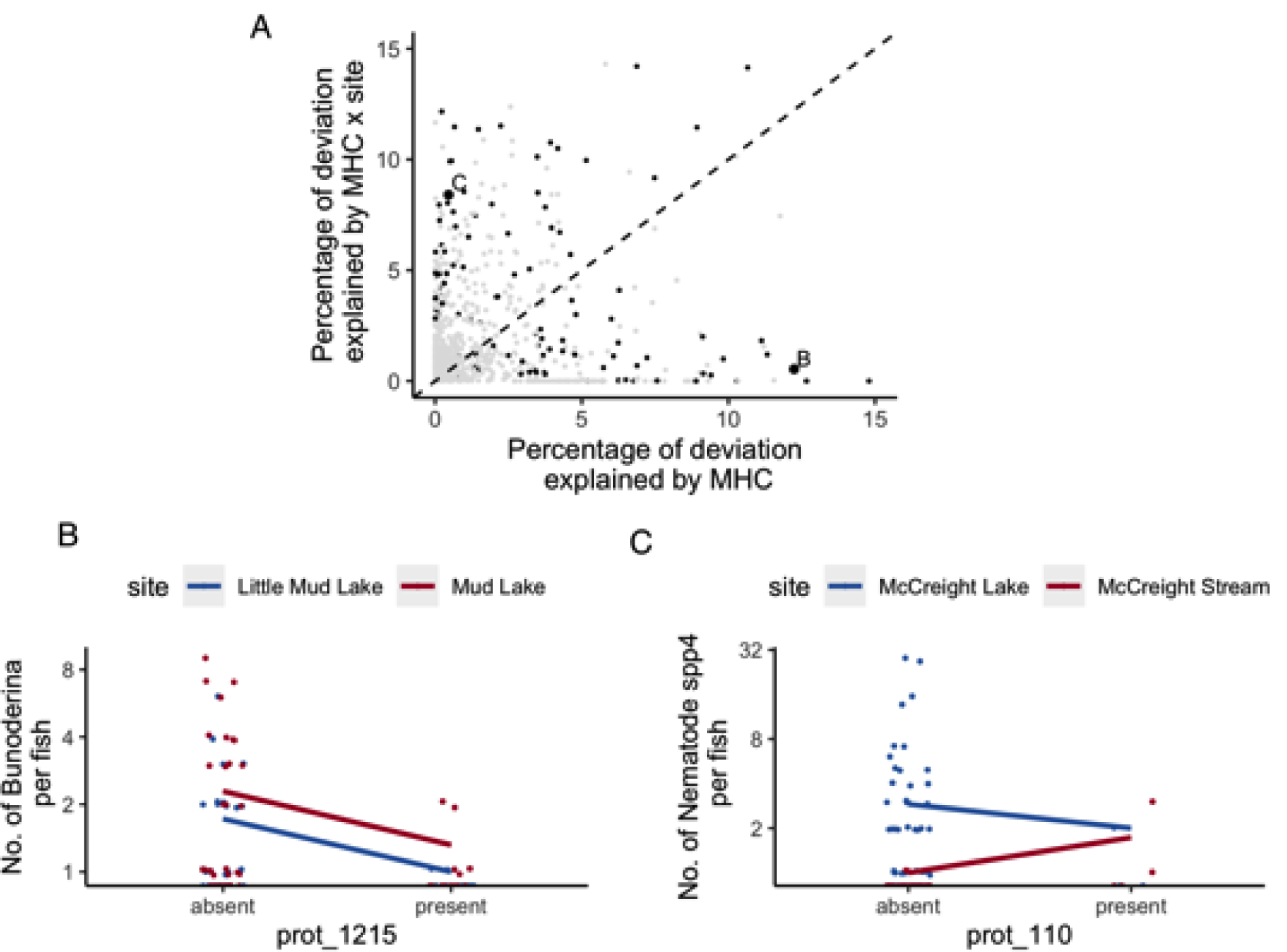
Testing predictions of frequency-dependent selection: inconsistent allele effects across sites. A. The percentage of deviation residuals explained by the MHC term and the MHC × sample site interaction term in the regression models, which regressed the infection load of certain parasite against a particular MHC allele, sample site, and the MHC × sample site interaction. Each dot represents a MHC-parasite combination. The dotted line is a line of equal effect size (slope of 1, intercept of 0). The models with either term significant are plotted with black dots, and all the non-significant models are in grey. Two significant examples were highlighted in A, and plotted in B and C. B. The effect of MHC allele prot_1215 on the infection load of parasite Bunoderina in Mud lake and Little Mud lake. C. The effect of MHC allele prot_110 on the infection load of parasite Nematode spp4 in McCreight lake and McCreight stream. Note that the y axes of B and C are on log scale.

#### c. Fluctuating selection

Despite spatial heterogeneity in the distribution of parasite species (Bolnick et al., 2019), we did not detect parasite-mediated spatially fluctuating selection. The Mantel test between the distance matrix of parasite community and the distance matrix of MHC alleles across all populations suggested that the two matrices were not correlated (r = 0.089, p=0.22, Fig. 5). Thus, although MHC alleles and parasite communities both differ strongly between lakes, there is no detectable relationship between them.

**Figure 5.**
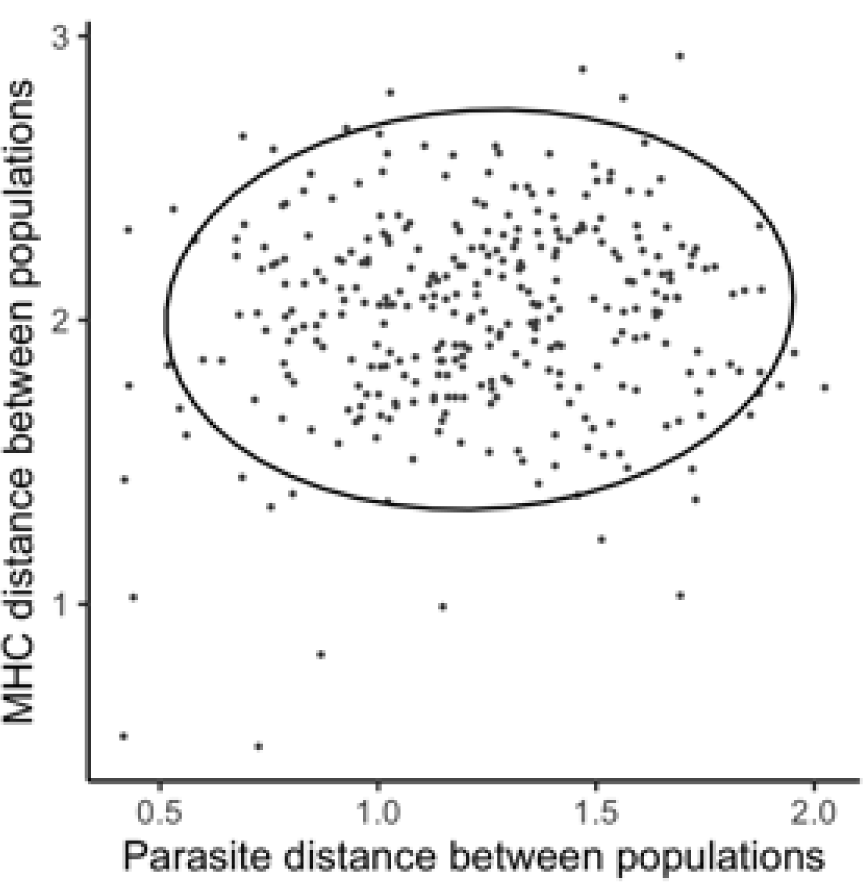
Testing predictions of spatially fluctuating selection. The parasite distance between populations is not detectably correlated with the MHC distance between populations. Each dot represents a pair of populations. The ellipse represents 95% confidence interval prediction.

#### d. Neutral evolution

We found that MHC diversity was significantly associated with genome-wide genetic diversity that should mostly reflect neutral processes. First, the average MHC allelic richness covaried positively with population mean heterozygosity (coefficient=25.4, F=6.57, df = 24, p = 0.02, Fig. 6A). Second, there was a strong positive correlation between MHC divergence between populations, and genome-wide divergence (Mantel test: r = 0.51, p=0.002, Fig. 6B). That is, more closely related populations tended to be more similar at their MHC loci as well.

**Figure 6.**
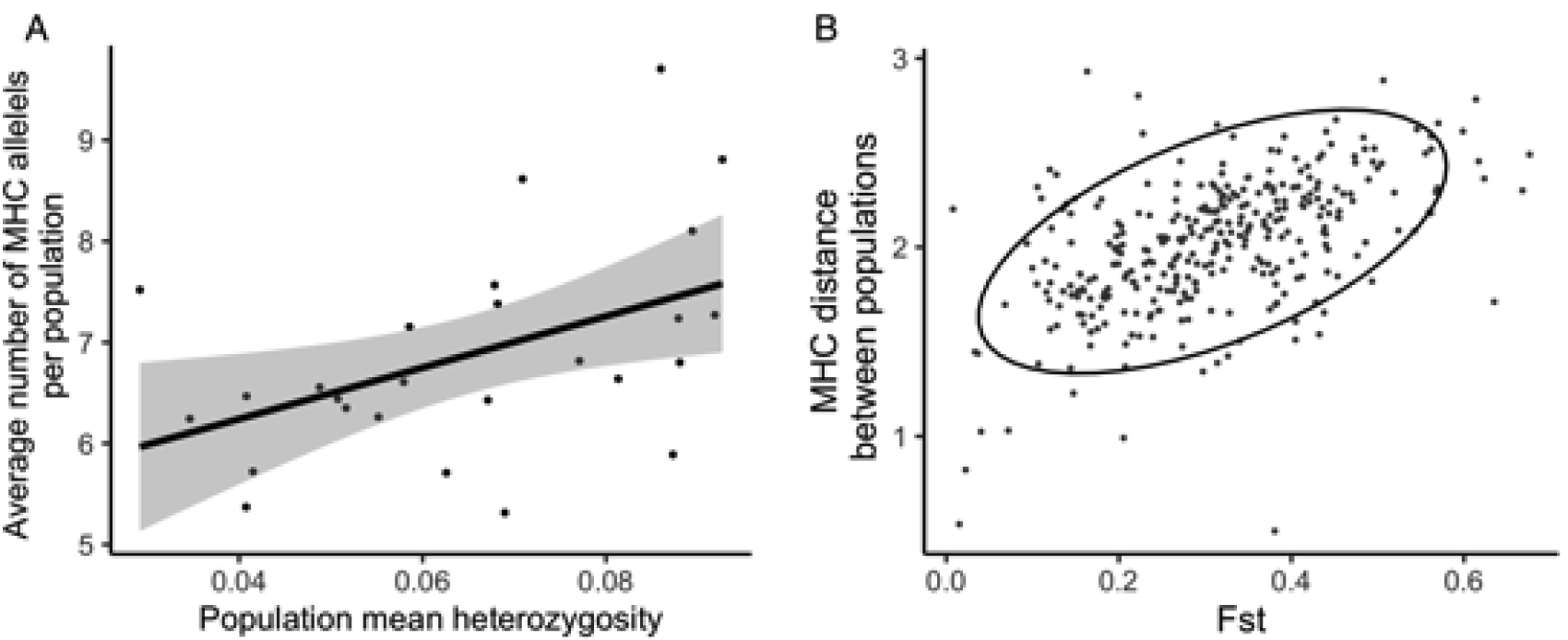
Testing predictions of neutral hypotheses. A. The average number of MHC alleles per population is positively associated with population mean heterozygosity. Each dot represents a population. The solid line represents linear regression prediction. Grey area indicates 95% confidence interval. B. The relationship between MHC distance between populations and Fst. Each dot represents a pair of populations. The ellipse indicates 95% confidence interval prediction.

## Discussion

This study represents an effort to comprehensively evaluate multiple major hypotheses concerning the maintenance of MHC diversity, using observational data from 26 stickleback populations. Despite the large scale of sampling, we did not find strong support for any of the three popular parasite-mediated selection hypotheses, namely heterozygote advantage, frequency-dependent selection, and fluctuating selection. In contrast, neutral processes seem to best explain MHC diversity in this system, both in terms of allelic richness and between-population divergence.

Consistent with previous studies in other vertebrates (Kaufman, 2018), MHC IIβ genes are highly polymorphic in the surveyed stickleback metapopulation. We found 1115 unique MHC IIβ alleles from 1277 individuals, with a majority of alleles only present in one population. However, although on average fish from lake habitats are infected with more diverse parasite species than the ones from river and estuary habitats (Bolnick et al., 2020), lake stickleback have less MHC allelic diversity than others. The extremely high level of polymorphism in MHC genes is often attributed to parasite-mediated selection; however, the discrepancy between parasite richness and MHC allelic diversity among habitat types suggests that parasite-mediated selection alone is unlikely to explain the high level of MHC polymorphism in the studied stickleback populations. This finding contradicts previous results from studies of stickleback inhabiting different habitats in Germany (K. M. Wegner, Reusch, & Kalbe, 2003), where lake fish had higher parasite diversity (as in this study), and higher MHC diversity (unlike this study).

The above finding is further supported through testing specific predictions derived from heterozygote advantage model. The heterozygote advantage hypothesis predicts that the number of MHC alleles in the host genome would be *maximized* to recognize a wide range of parasites. But previous studies (K. Mathias Wegner et al., 2003) in stickleback found support for the theoretical model that the number of MHC alleles should be *optimized* at an intermediate level, rather than *maximized*. The argument is that too many MHC alleles would reduce the T cell portfolio due to negative selection of self-recognition T cell receptors. In this study, we applied similar analytical approaches as Wegner et al. (2003) to a larger sample size and many more populations. Contrary to those previous results, we did not find support for either the *maximizing* or *optimizing* hypothesis.

Negative frequency-dependent selection suggests that the frequency of MHC alleles might experience a cyclical pattern through time, with different low frequency alleles playing the role of conferring resistance at different time points. For negative frequency dependence to be effective, on average, it must be true that currently rare alleles exhibit an advantage over common ones. In this study, we found that allele prevalence in general is not associated with whether an allele is effective in parasite resistance. We further examined whether alleles have consistent effects across sites. We found inconsistent allele effects in some cases, but we also found consistent effects in a larger fraction of cases. So negative frequency-dependent selection might be able to explain a limited number of allele-parasite dynamics in our study. But, in most cases MHC alleles confer equivalent protection in multiple isolated populations, inconsistent with cyclic changes due to coevolution. Ideally, to demonstrate negative frequency-dependent selection, one would need to carry out a longitudinal study that samples parasite genotypes and host genotypes through time. However, this type of study is very challenging to implement in a field setting, especially for annually reproducing animals.

Finally, fluctuating selection, either temporally or spatially, is frequently invoked to explain the maintenance of genetic diversity across a landscape (Charbonnel & Pemberton, 2005; Schemske & Bierzychudek, 2007). Our sampling sites on Vancouver Island differ in many physical parameters, such as water area, depth, temperature, flow speed, *etc*. It is also documented that parasite communities differ significantly among different sites (Bolnick et al., 2020). This spatial heterogeneity in parasite communities is due to abiotic and biotic variables (e.g., lake size, fish diet; Bolnick et al. (2019)), and could generate spatially fluctuating selection that maintains MHC diversity. However, we did not find significant correlation between the parasite distance matrix and the MHC distance matrix, suggesting that the spatial heterogeneity in parasite community could not explain the observed spatial variation in MHC diversity.

Interestingly, we found that neutral processes could contribute to MHC diversity in two different analyses: (a). populations with higher genomic heterozygosity also have higher MHC allelic richness, and (b). less divergent populations (low genomic Fst) have similar MHC genotypes and frequencies. Similar to analysis (a), a previous study in stickleback (K. M. Wegner et al., 2003) also found a positive, albeit weak, association between neutral genetic diversity and population MHC allelic diversity. The role of neutral processes in the maintenance of MHC diversity has often been studied in the context of conservation, which is usually concerned with small and isolated populations (Miller & Lambert, 2004; Seddon & Ellegren, 2004). Our result suggested that at larger scales, neutral processes are also important factors in explaining MHC diversity within- and between-population, even when there is no specific evidence for past bottlenecks.

Parasite-mediated selection is believed to have an important role in maintaining MHC polymorphism. Consistent with this, we found some associations between certain parasites and individual MHC allele. However, we did not find strong support for parasite-mediated selection hypotheses in the scale of metapopulations. There are several possible explanations. First, in this study, we examined the three hypotheses separately, but in reality, the mechanisms are not mutually exclusive. For example, low frequency alleles are more likely to be in a heterozygous state in a population. Therefore, if an allele were effective in resistance at low frequency, it would be consistent with both negative frequency-dependent selection and heterozygote advantage. Furthermore, if these mechanisms work in combination in different MHC-parasite pairs or in different populations, we may not detect a clear signal of each mechanism when combining data from multiple MHC-parasite pairs and many populations. Second, parasites are not the only selective agents that act upon MHC loci. MHC genes are also involved in interactions with other species, such as symbionts comprising the gut microbiome (Bolnick et al., 2014), and within species interactions, such as mate choice (Milinski et al., 2005). It is unknown which MHC genes are relevant to which selective forces. If other selective forces are also at work, the influences of parasite-mediated selection on MHC diversity could be obscured. Third, we only measured parasite richness and MHC allelic richness in this study to represent parasite diversity and MHC diversity, respectively, so the detailed differences among individual parasite and individual MHC alleles were not considered. In the future, it is worthwhile to test the parasite-mediated selection hypotheses with diversity measurements that take into account of nuances such as the phylogenetic relationship of parasite taxa and the sequence divergence between distinct MHC alleles. Finally, our survey does not span multiple seasons, so we cannot evaluate how MHC variation is shaped across time by temporally fluctuating selection.

To summarize, we used a large-scale field survey to evaluate the three popular parasite-mediated selection hypotheses on MHC diversity, namely, heterozygote advantage, negative frequency-dependent selection, and fluctuating selection. We found that neutral processes best explain MHC diversity (allelic richness and population divergence), instead of those parasite-mediated selection mechanisms. Because MHC diversity can be influenced by many selective forces, and the outcome of this selection could be further shaped by spatial and temporal heterogeneity, our study suggests it may not be possible to parse out how each selective force influences MHC diversity at large scales directly from observational data. We propose that it is worthwhile to instead investigate how each selective force acts upon MHC diversity with experimental approaches (*e*.*g*. controlling other selective forces) at smaller scales. Combining insights from these small-scale controlled studies in a step-wise manner can be a fertile future direction to understand the complex process of MHC evolution and diversification.

## Acknowledgment

This work was supported by the David and Lucille Packard Fellowship for Science and Engineering (DIB), the Howard Hughes Medical Institute (Early Career Scientist award to DIB), and by NIH NIAID grant 1R01AI123659-01A1. Field sampling in 2009 was made possible by assistance from Todasporn Rodbumrung, Chris Harrison, Will Shim, Yuexin Jiang, Will Stutz. Will Shim provided parasite information in Table S2.

## Data Accessibility

MHC allelic diversity and parasite species richness data will be deposited in Dryad. Sample site and parasite taxa information is included as supplemental materials. The code used to perform data analysis is available in the GitHub repository: https://github.com/foenpeng/MHC_analysis

## Author Contribution

DIB conceived and designed this project with Will Stutz. DIB and WS performed the field survey. KB carried out the dissections. SHW performed the MHC sequencing, SDH performed genotype scoring. FP performed data analysis and wrote the manuscript with input from DIB. All authors contributed in the revision of the manuscript.

## Notes

### Competing Interest Statement

The authors have declared no competing interest.

### Summary of Updates

coauthor's email and some formatting

