## Supplementary material for "What evolutionary processes maintain MHCIIβ diversity within and among populations of stickleback?": Table S1

**Supplemental materials**

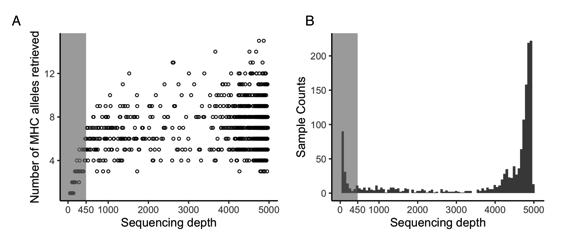

Figure S1. A. The relationship between sequencing depth and the number of MHC alleles. Each dot represents a fish sample. B. The distribution of sequencing depth among samples. Individuals with sequencing depth < 450 were excluded from analysis.

Table S1. Sample site information

| site_name | site_ID | habitat_type | watershed | sample_size |
| --- | --- | --- | --- | --- |
| Blackwater Lake | BLK | Lake | Amor River | 58 |
| Bob Lake | BOB | Lake | Pye River | 25 |
| Brewster Lake | BRW | Lake | Campbell River | 42 |
| Cecil Lake | CEC | Lake | Amor River | 63 |
| Cedar Lake | Ced | Lake | Amor River | 65 |
| Cranberry Lake | CRB | Lake | Mohun River | 45 |
| Campbell River Marsh | CRM | Estuary | Campbell River | 60 |
| Echo Lake | ECH | Lake | Quinsam River | 45 |
| Fry Lake | FRI | Lake | Campbell River | 59 |
| Gray Lake | GRL | Lake | Campbell River | 60 |
| Higgens Lake | HIG | Lake | Campbell River | 54 |
| Lawson Lake | LAW | Lake | Campbell River | 78 |
| Lower Campbell Lake | LCA | Lake | Campbell River | 18 |
| Little Mud Lake | LM | Lake | Amor River | 51 |
| McCreight Lake | MCR | Lake | Amor River | 47 |
| McCreight Stream | MCS | Stream | Amor River | 34 |
| Merril Lake | MER | Lake | Campbell River | 25 |
| Mohun Lake | MOH | Lake | Mohun River | 53 |
| Mud Lake | BM | Lake | Amor River | 52 |
| Muskeg Lake | MUS | Lake | Amor River | 51 |
| Pye Creek | PYS | Stream | Pye River | 52 |
| Pye Outlet | PYO | Estuary | Pye River | 59 |
| Sayward Estuary | SAY | Estuary | Salmon River | 59 |
| Stella Lake | STE | Lake | Pye River | 49 |
| Upper Campbell Lake | UPC | Lake | Campbell River | 55 |
| Village Bay Lake | VBL | Lake | Village Bay | 18 |

Table S2. Parasite information

| Name | Final_hosts | Intermediate_hosts_before_fish | Mode_of_transmission | Notes&references |
| --- | --- | --- | --- | --- |
| Acanthocephanus | birds | crustaceans | oral | https://journals.plos.org/plosone/article?id=10.1371/journal.pone.0028285 |
| Anisakis | birds or mammals | crustaceans | oral | genus of nematode. Phylogeny and more info here and references therein: https://pubmed.ncbi.nlm.nih.gov/16800118/ |
| Argulus | stickles depending on the sp | n.a. | direct attachment | genus of crustacean, single host, direct transmission from other stickles. They're called marine lice because they work just like ones |
| Blackspot | birds probably | snails | direct penetraction and attachment | blackspots are cysts of mostly trematodes in the skins of fish. They get transmitted to final hosts when stickles are ingested. Hard to tell what spp they are from cysts. |
| Bunodera | stickle | crustacean? | oral | https://link.springer.com/article/10.1007%2Fs00436-018-5858-y |
| Bunoderina | stickle | crustacean? | oral | https://www.nrcresearchpress.com/doi/abs/10.1139/cjr36d-003#.XzStLS2z0Wo |
| Capillaria | stickle? | environement or crustaceans? | oral | see references here: https://en.wikipedia.org/wiki/Capillaria_(nematode) |
| Cestode spp2 | birds or fish | crustaceans or insect larva | oral? |  |
| Cestode spp3 | birds or fish | crustaceans or insect larva | oral? |  |
| Cestode spp4 | birds or fish | crustaceans or insect larva | oral? |  |
| Cestode spp5 | birds or fish | crustaceans or insect larva | oral? |  |
| Crepidostomum | stickle | crustacean? | oral | https://pubmed.ncbi.nlm.nih.gov/17436963/ |
| Cystidicola | bigger fish? | environement or crustaceans? | oral? | https://www.researchgate.net/publication/237178203_A_Revision_of_the_Genus_Cystidicola_Fischer_1798_Nematoda_Spiruroidea_of_the_Swim_Bladder_of_Fishes and https://pubmed.ncbi.nlm.nih.gov/29906216/ |
| Diplostomum spathaceum | birds | snails | direct penetraction and attachment | https://www.researchgate.net/figure/Phylogenetic-position-of-the-genus-Diplostomum-estimated-under-the-maximum-likelihood_fig1_278791759 |
| Dermocystidium | stickles probably | n.a. | probably skin penetration | thought to be related to fungi. http://www1.nencki.gov.pl/pdf/ap/ap718.pdf Also, look for https://www.nature.com/articles/196958a0 |
| Ergasilus | stickle | n.a. | direct attachment | https://www.researchgate.net/publication/5904390_Phylogeny_of_freshwater_parasitic_copepods_in_the_Ergasilidae_Copepoda_Poecilostomatoida_based_on_18S_and_28S_rDNA_sequences |
| Eustrongyloides | birds | polycheatas | oral | https://pubmed.ncbi.nlm.nih.gov/22924908/ |
| Glugea | stickle | n.a. | oral, mother to offsrping thru eggs | Microsporidian: https://en.wikipedia.org/wiki/Microsporidia |
| Nematode spp2 | | crustaceans or insect larva | oral? |  |
| Nematode spp3 | | crustaceans or insect larva | oral? |  |
| Nematode spp4 | | crustaceans or insect larva | oral? |  |
| Nematode spp6 | | crustaceans or insect larva | oral? |  |
| Nematode spp7 | | crustaceans or insect larva | oral? |  |
| Nematode spp10 | | crustaceans or insect larva | oral? |  |
| Neoechinorhyncus | birds | crustacean | oral | https://pubmed.ncbi.nlm.nih.gov/24064255/ |
| Neoechinocephalus | birds? | crustacean? | oral? | Seems to be the same family as Neoechinorhyncus above. https://pubmed.ncbi.nlm.nih.gov/24064255/ |
| Proteocephalus | bigger fish? | crustacean | oral | https://www.sciencedirect.com/science/article/abs/pii/S0020751901002260 |
| Proteocephalus spp2 | bigger fish? | crustacean | oral | https://www.sciencedirect.com/science/article/abs/pii/S0020751901002260 |
| Raphidascaris | bigger fish? | crustacean or insect larva | oral | https://www.researchgate.net/publication/329843388_A_new_species_of_Raphidascaris_Nematoda_Raphidascarididae_infecting_the_fish_Gymnogeophagus_balzanii_Cichlidae_from_the_Pantanal_wetlands_Brazil_and_a_taxonomic_update_of_the_subgenera_of_Raphidascari |
| Schistocephalus | birds | crustaceans | oral | https://journals.plos.org/plosone/article?id=10.1371/journal.pone.0022505 |
| Thersitina | stickle | n.a. | direct attachment | https://www.gbif.org/species/2110447, https://link.springer.com/article/10.1023/B:SYPA.0000010683.46287.65 , |
| Unionidae | free living adults | n.a. | direct attachment | Family of freshwater clams where larvae are parasitic or attach to fish for dispersal. https://www.sciencedirect.com/science/article/abs/pii/S1055790316302202 |
| Diplostomum spp | birds | snails | direct penetraction and attachment | https://www.researchgate.net/figure/Phylogenetic-position-of-the-genus-Diplostomum-estimated-under-the-maximum-likelihood_fig1_278791759 |

Table S3. Model selection for heterozygote advantage analysis

| **Model** | **Degree of freedom** | **AIC** |
| --- | --- | --- |
| Parasite.richness ~ log_std_length + (1\|site_name) | 3 | 4226.024 |
| Parasite.richness ~ log_std_length + x + (1\|site_name) | 4 | 4226.531 |
| Parasite.richness ~ log_std_length + sqx + (1\|site_name) | 4 | 4226.597 |
| Parasite.richness ~ log_std_length + x + (1\|site_name) + (x\|site_name) | 7 | 4231.633 |
| Parasite.richness ~ log_std_length + sqx + (1\|site_name) + (sqx\|site_name) | 7 | 4231.724 |
| Parasite.richness ~ log_std_length + x + sqx + (1\|site_name) + (x\|site_name) | 8 | 4233.632 |
| Parasite.richness ~ log_std_length + x + (1\|site_name) + (x\|site_name) + (sqx\|site_name) | 10 | 4237.633 |
| Parasite.richness ~ log_std_length + x + sqx + (1\|site_name) + (x\|site_name) + (sqx\|site_name) | 11 | 4239.632 |
